# Investigation of V1016G *kdr* Mutation during Pyrethroid Insecticide Resistance in *Aedes aegypti* from Nagapattinam district of Tamil Nadu, India

**DOI:** 10.1101/2025.08.05.668611

**Authors:** Rajalakshmi Anbalagan, Priyadharshini, Mortein Henri, Farhat Syed Kaseem, Sathya Jeevitha, Mansi Malik, Balachandar Vellingiri, Anand Setty Balakrishnan, Sulochana Shekhar, Jayalakshmi Krishnan

**Affiliations:** Department of Biotechnology, School of Integrative Biology, Central University of Tamil Nadu, Thiruvarur, 610 001, Tamil Nadu,India; Tata Institute for Genetics and Society, instembuilding, Firstfloor, NCBS-GKVK campus, Bengaluru, Karnataka,560065,India; Human Cytogenetics and Stem Cell Laboratory, Department of Zoology, School of Basic Sciences, Central University of Punjab, Bathinda, Punjab 151401, India; Department of Epidemiology and Public Health, Central University of Tamil Nadu, Thiruvarur, 610 001, Tamil Nadu, India; Central University of Tamil Nadu, Thiruvarur, 610 001, Tamil Nadu, India

**Author notes:** Corresponding author, Department of Biotechnology, School of Integrative Biology, Central University of Tamil Nadu, Thiruvarur, 610 001, Tamil Nadu, India.

**Keywords:** *Aedes aegypti*, allele specific PCR, Insecticide susceptibility, heterozygous mutation, Nagapattinam, V1016G *kdr* mutation

## Abstract

**Background:** *Aedes aegypti* is a primary vector of Dengue, Chikungunya, and Zika viruses, and its control relies heavily on pyrethroid insecticides. However, the development of resistance in mosquito populations poses a major challenge to effective vector control. This study aimed to investigate the presence of the V1016G knockdown resistance (*kdr*) mutation in *Aedes aegypti* collected from Nagapattinam district, Tamil Nadu, India. Where insecticide uses is common.

**Methodology:** *Aedes* immature were collected from selected field sites and subjected to insecticide susceptibility testing using World Health Organization (WHO) guidelines. Genomic DNA was extracted, and allele-specific PCR was performed to detect the V1016G mutation in the voltage-gated sodium channel gene *(vgsc)*.

**Principal findings:** The analysis revealed the presence of the V1016G mutation in heterozygous form (VG) among the tested samples. No homozygous mutants (GG) were detected. The presence of heterozygous individuals indicates an early stage of resistance development, likely due to ongoing insecticide selection pressure. Bioassay results showed reduced susceptibility to pyrethroids in these populations, supporting the molecular findings.

**Conclusion:** This study provides baseline data on the genetic mechanism of pyrethroid resistance in *Aedes aegypti* from Nagapattinam district. Regular monitoring of *kdr* mutations is essential to detect resistance early and guide insecticide use policies. Integrating molecular surveillance with conventional control measures will help delay the spread of resistance and improve the long-term success of vector management programs. The findings highlight the need for alternative control strategies and responsible insecticide use to maintain the effectiveness of current interventions.

## Introduction

Mosquitoes are the major epidemiological concern due to their critical role in the transmission of deadly arboviral diseases such as Dengue Virus (DENV), Zika virus, Yellow fever and Chikungunya [1]. Among the various vectors, *Aedes aegypti*, is the principle vector responsible for the spread DENV [2]. DENV stand out as a global threat to humans that continue to endanger millions of live each year. In 2023, WHO ranked dengue as the 3^rd^ emerging disease, following a surge in severe outbreaks across several countries [3]. As on 30^th^ April 2024 moreover 7.6 million dengue cases have been reported including 3.4 million confirmed cases includes 16,000 severe cases with 3000 deaths [4]. This vector is morphologically similar to *Aedes albopictus*, with distinctive black and white markings on their back and legs [5]. However, it uniquely exhibits strong anthrophilic behavior [6]. This adaptability brings it into close proximity with human dwellings and enables it to exploit both natural and artificial containers as breeding sites [7-8]. with its global spread often facilitated by global transportation [9].

Vector control programme have implemented various control measurements to mitigate the risk of disease transmission, such as the use of insecticides, indoor residual sprayings (IRS), tropical repellents, and insecticide-treated bed nets respectively. Additionally, biological agents [10] and innovative technologies such as nanoparticles have also been employed to eliminate mosquito breeding sites [11]. Despite these extensive approaches, vector control strategies still rely mainly on chemical insecticides, including pyrethroids, organophosphates and carbamates [12]. Pyrethroid insecticides act as neurotoxins by targeting the voltage sensitive sodium channel (VSSC). These channels in insects are significantly more susceptible to pyrethroids compared to those mammals [13]. Pyrethroids work by destructing the functions of voltage-sensitive sodium channels, keeping them open for longer periods and interfering with normal nerve signaling [14]. Resistance towards pyrethroid occurs through two major mechanisms: the first mechanism comprises metabolic or enzymatic resistance, and the second mechanism is knockdown resistance, or *kdr* [15]. The hallmark of pyrethroid effects on insects is their ability to cause “knockdown,” a condition marked by rapid paralysis. This occurs because pyrethroids keep sodium channels in an activated state for extended periods, disrupting the transmission of nerve signals. Knockdown resistance, often abbreviated as *kdr*, arises primarily from mutations in the sodium channels that reduce this sensitivity [13]. Researchers have identified over 50 mutations in sodium channels linked to pyrethroid resistance in insect pests and disease vectors affecting humans. Many of these mutations have been linked to pyrethroid resistance in *Aedes aegypti* [16]. Many studies have identified various *kdr* mutation, including G923V, L982W, I1011M/V, S989P, V1016G/I, F1534C, and D1763Y [17] among these V1016G, V1016I, and F1534C were consistently identified in insecticide-resistant populations [13]. The amino acid substitutions V1016G and F1534C are the most common and well-established mutations driving pyrethroid resistance in *Aedes aegypti* mosquitoes [18]. In this present study, we focused on insecticide resistance status in *Aedes aegypti* against pyrethroids and investigated the V1016G *kdr* mutation in *Aedes aegypti* populations, which has not been extensively explored in Tamil Nadu, particularly in the Nagapattinam region—an area known for recurring arboviral disease outbreaks. This research will provide insights into the mechanisms of insecticide resistance in the local mosquito population and contribute to improved vector control strategies.

## Materials and methods

### Field collection of *Aedes aegypti* from Nagapattinam

Larvae and pupae samples of *Aedes aegypti* were from Nagapattinam district during monsoon and postmonsoon seasons (November to February, 2025). Nagapattinam District is one of the 38 districts of Tamil Nadu state in southern India. Nagapattinam District is located along the Bay of Bengal and is geographically positioned at 10.7906°N latitude and 79.8428°E longitude. Totally Nagapattinam has 11 blocks (https://www.nagapattinam.nic.in/). The district is stretching across eleven blocks namely keelaiyur, Kilvelur, Kollidam, Kuttalam, Mayiladuthurai, Nagapattinam, Sembanar koil, Sirkazhi, Talanayar, Thirumarugal, and Vedaranniyam. The survey was conducted across various locations by means of random sampling from these eleven blocks, where dengue cases were highly reported. The Global Positioning System (GPS) locations of samples collected areas were mapped using the software, ArcGIS 10.8, from the department of geography, CUTN as mentioned in the Figure 1.

**Figure 1:**
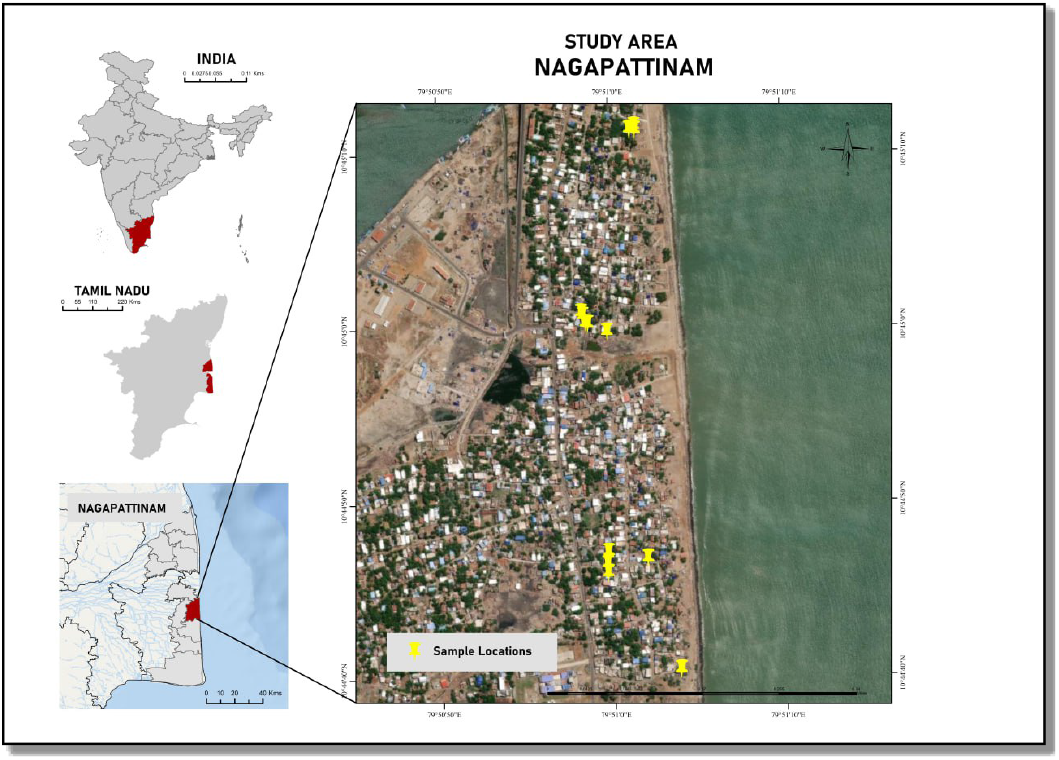
Map showing locations of sample collected areas-Nagapattinam district (Source: ArcGIS 10.8)

The immature samples were collected from the different key breeding sites of *Aedes aegypti* by using the Standard methods of WHO. Further, sampling was done by using standard white enameled dipper (250 ml) which helps to see the mosquitoes in deep water containers (WHO, 2009). Further, a dipper was dipped into the water logging at angle of about 45°. Wide mouth glass pipette and a dropper were used to separate larvae and pupae (WHO, 2009).

### Laboratory rearing of Mosquito

Immature samples were initially collected in separate plastic containers from different breeding sites and then transported to the Vector Biology Research Laboratory (VBRL), Department of Biotechnology, CUTN. The collected larvae were separated to the different enamel trays and maintained in the VBRL by providing larval food (dog biscuit and yeast powder mixed in the ratio 3:1) and the emerged adult mosquitoes were identified using standard morphological keys. Adult mosquitoes were maintained on 10% glucose solution provided via cotton pads soaked with resin. Female mosquitoes were fed with chicken blood through a membrane feeder to facilitate their egg laying. Subsequently, Ovitraps were set up to collect eggs from the gravid females. These simple containers, designed to attract female mosquitoes for oviposition, were strategically placed and contained water along with substrates such as country man No.1 filter paper to serve as egg-laying surfaces. The deposited eggs were removed, dried, and stored at room temperature for further use. F1 generation of *Aedes aegypti* mosquitoes were used for insecticide susceptibility test.

### Insecticide susceptibility test (WHO Tube test)

Adult insecticide susceptibility tests were performed according to the WHO susceptibility test kit method. Mosquitoes were exposed to pyrethroids insecticides namely Deltamethrin 0.05% and permethrin 0.75%, six clean white paper sheets were rolled into a green-dot-marked holding tube and secured with silver clips. Female *Aedes aegypti* mosquitoes (F1 generation) were aspirated and transferred to the tube. Batches of 25 mosquitoes were acclimatized for one hour in the holding tube before being exposed to insecticide impregnated papers in red-dot-marked exposure tubes. Four test replicates and two controls (with yellow-dot-marked tubes and oil-impregnated papers such as OP-control and PY-control) were used, each containing 25 mosquitoes. After one hour of exposure, mosquitoes were returned to holding tubes with sugar-soaked cotton pads for 24 hours Knockdown rates were recorded at 10-minute intervals during exposure, and 24-hour mortality was calculated using the below mentioned formula. Observed resistant mosquitoes were stored at -80°C for further process.

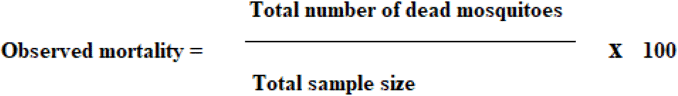

Abbott’s formula **(Abbott, 1925)** was not applied for the calculation of corrected mortality, as no mortality was observed in the control group. The resistance status of mosquitoes was determined according to WHO guidelines. A mortality rate of 98–100% indicates susceptibility (S), 90–97% suggests possible resistance (PR), and a mortality rate below 90% confirms resistance (R) (WHO, 2022)

### Molecular characterization

#### Isolation of genomic DNA from mosquito sample

Totally, 81 individuals were identified as resistant based on the bioassay results. The resistant mosquitoes were selected for further molecular analysis. DNA samples were isolated from permethrin and Deltamethrin resistant samples using the commercial DNeasy Blood and Tissue (Qiagen, Germany) kit following manufacturer protocol and DNA concentration was quantified using an Eppendorf Biospectrometer based on absorbance at 260nm.Mosquito was processed individually for Allele-Specific PCR (AS-PCR) to detect the presence of the V1016G *kdr* mutation. **AS-PCR for V1016G *kdr* mutation**

The point mutation Valine (V) to Glycine (G) in the voltage gated sodium channel at V1016G residue (*kdr*) was identified by Allele specific PCR (AS-PCR). Following the method developed by [31] with minor modification. All the PCR requirements were retrieved from -20°C and kept on the icebox prior to use. Each reaction was performed in a 25 μl volume consisting of 12.5 μl of Zensus 2X Taq DNA PCR mastermix, 1 μl of common forward primer, 0.95 μl of each reverse primer specific for either Glycine or Valine, 2 μl of template DNA and finally 8.55 μl nuclease free water, the volume of water is adjusted based on the quantity of DNA, primer details have given in the Table 1. The PCR amplification was initiated with an initial DNA denaturation step at for 94°C for 2 minutes,, followed by 35 cycles containing of denaturation at 94°C for 30 seconds, annealing at 58.5°C for 30 seconds, and extension at 72°C for 30 seconds minutes. A final extension was carried out at 72°C for 10 minutes to complete the reaction. Due to the inclusion of GC-rich tails of different lengths in the primers, the amplified products were distinguishable by size—60 bp for the Valine allele and 80 bp for the Glycine allele. Further PCR amplification products were loaded onto a 2% Agarose gel and run for 1hour at 75V in TBE buffer.

**Table 1:**
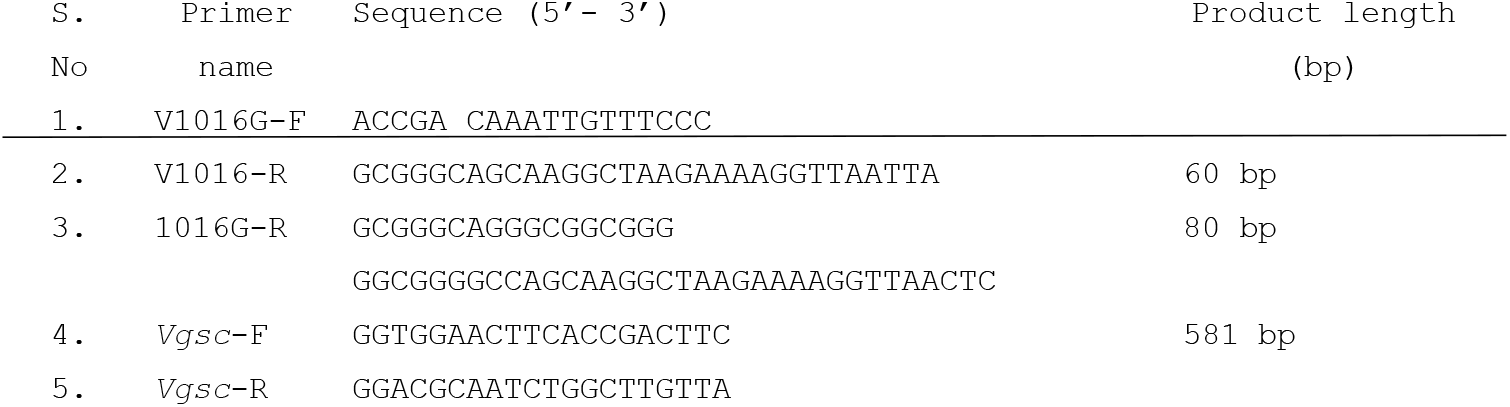
Primer sequences for AS-PCR and PCRp (Stenhouse *et al*., 2013)

#### Target gene-amplification PCR

To confirm the AS-PCR results, conventional PCR was performed to amplify the region of domain II subunit 6 in the *vgsc* gene encompassing the V1016G mutation. Each reaction was performed in 20 µl volume consisting of 10 µl Zensus 2X Taq DNA PCR master mix, 0.4 µl of forward primer and 0.4 µl of reverse primer, 7 µl of amplified product of AS-PCR, and finally 2.2 µl of nuclease free water,. PCR cycling condition were initiated with initial denaturation at 95°C for 2 minutes, followed by 35 cycles consisting of denaturation at 95°C for 30 seconds, annealing at 63°C for 30 seconds, denaturation at 72°C for 30 seconds and at last final extension at 72°C for 2mins to complete the reactions.

## Results

### Investigation of Resistance Patterns to Insecticides in *Aedes aegypti*

Extensive *Aedes* mosquitoes were collected from the various blocks of Nagapattinam, and tested for resistant to pyrethroids Table 2. The results indicated that *Aedes aegypti* was possible resistance to Deltamethrin (0.05%) with a recorded mortality rate of 96% whereas, the insecticide susceptibility tests for Permethrin (0.75%) indicates the confirmed resistance status with mortality rate of 84% as shown Figure 2.

**Table 2:** Insecticide susceptibility status (n = number of samples)

| S.No | Insecticides | No. of mosquitoes exposed<br>( Replicates n) | Mortality% | Status |
| --- | --- | --- | --- | --- |
| 1. | Deltamethrin (0.05%) | 150 | 96 | Possible resistance |
| 2. | Permethrin (0.75%) | 150 | 84 | Resistant |

**Figure 2:**
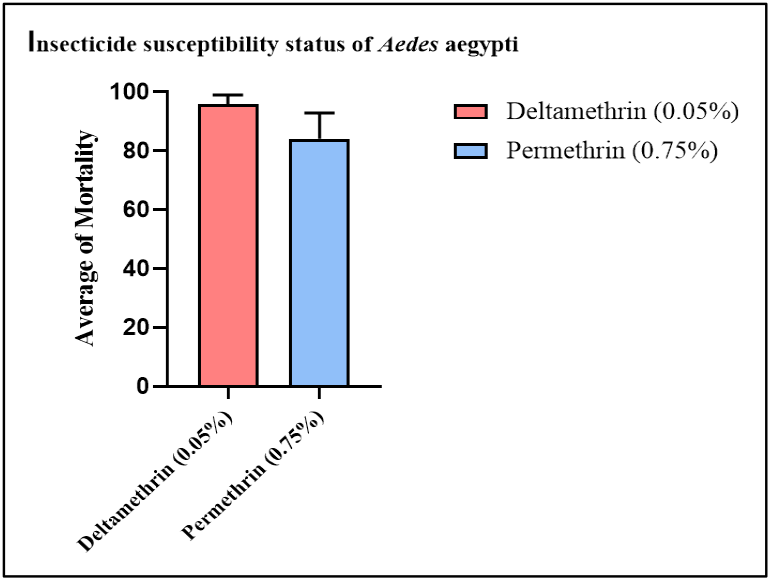
insecticide susceptibility status of *Aedes aegypti* from Nagapattinam.

Further, the resistant mosquito samples from the insecticide bioassay were subjected to molecular analysis to detect the V1016G *kdr* mutation using AS-PCR. The results revealed that all resistant samples carried the heterozgyous mutation (V/G) in the resistant individuals V1016G mutation, with no genotypic variation observed among them. Consequently, correlation analysis between genotype and phenotype could not be performed. To validate the AS-PCR results, the same samples were used for the amplification of the Voltage-gated sodium channel (*vgsc*) gene, specifically domain II-segment 6. Since the amplified product sizes (60bp for valine and 80bp for glycine) are too close to distinguish clearly in single-tube reaction as shown in Figure 3, a double-tube reaction approach was also used for each sample to improve band resolution and clarity as shown in Figure 4 and Figure 5. Later the band was visualized on 2% Agarose gel with the help of gel documentation system (BioBase).

**Figure 3:**
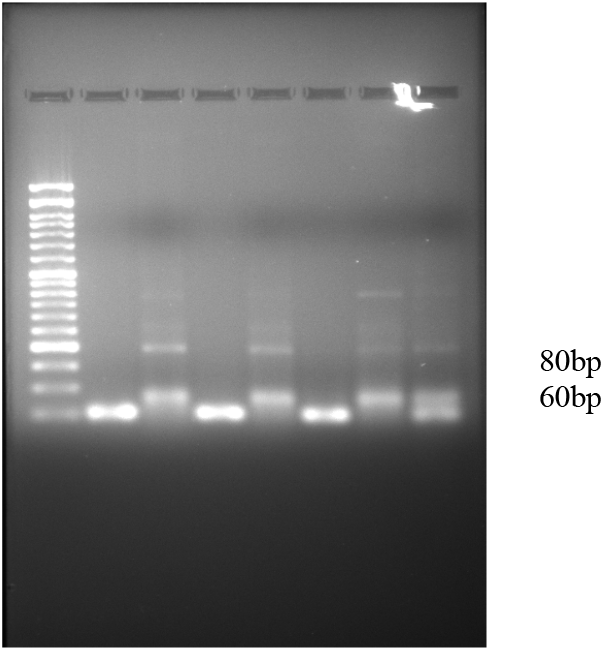
Gel image of deltamethrin-resistant samples. Lane 1: 50bp DNA ladder lane 2-7: double tube reaction is performed for resistant sample (60 bp valine wild type; 80bp glycine for mutant type) lane 8: single tube reaction was performed.

**Figure 4:**
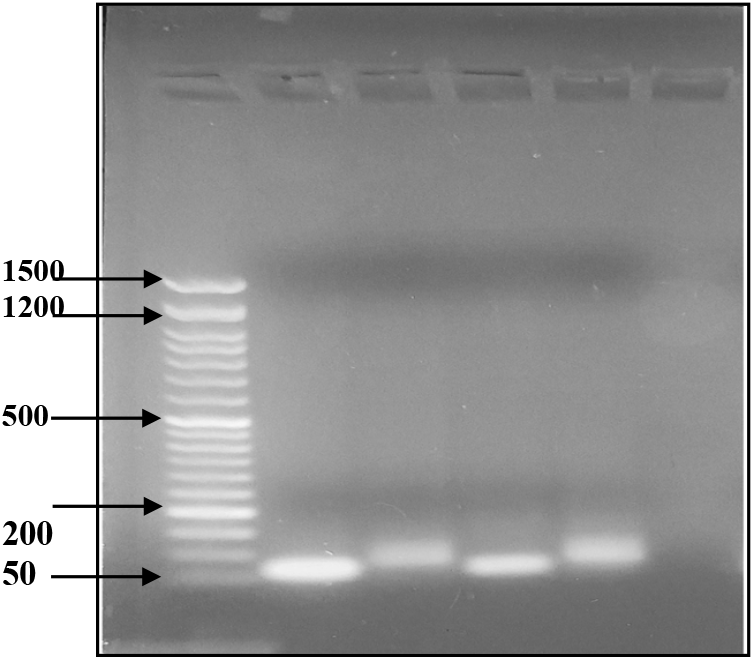
Gel image of Permethrin resistant samples. Lane 1: 50bp DNA ladder lane 2-5: double reaction was performed for samples lane 6: Negative control.

**Figure 5:**
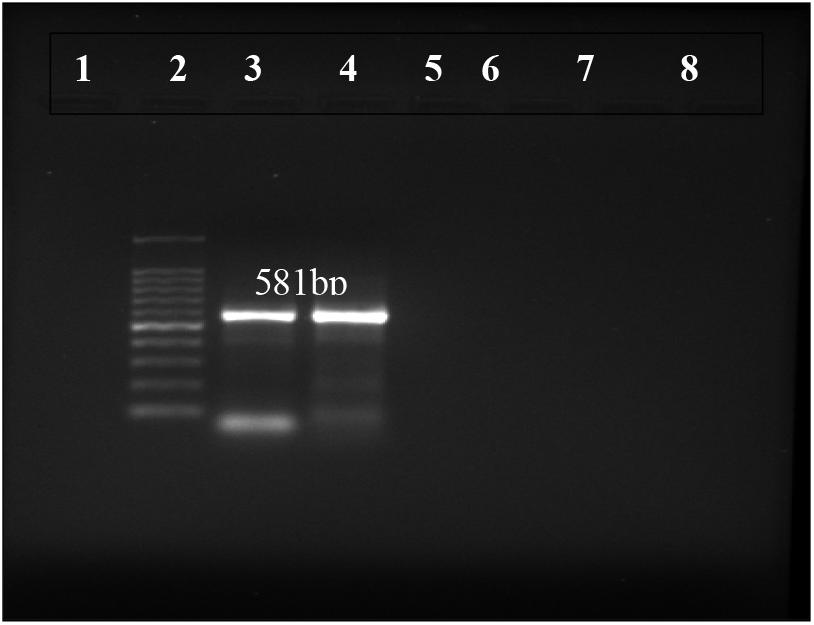
PCR amplification of AS-PCR amplified product using primers specific for the *vgsc* gene. **Lane 2**: 50bp DNA ladder, **Lane 3&4**: AS-PCR amplified Product of product size of **581bp**

## Discussion

*Aedes aegypti* is a major vector responsible for the transmission of Dengue worldwide. To mitigate the transmission several control measurements have been established such as usage of insecticides such as organophosphates, pyrethroids etc. Although insecticides have been extensively used for mosquito control, the emergence of resistance towards various insecticides has made the control strategies ineffective [19]. As discussed in the review of literature, one of the key mechanisms behind this resistance is the presence of knockdown resistance (*kdr*) mutation in Voltage-gated sodium channel *vgsc* gene [20] which reduces the efficacy of pyrethroids. Therefore the present study, was conducted to assess the insecticide susceptibility status of *Aedes aegypti* population from various location of Nagapattinam District and also detected the presence of V1016G *kdr* mutation which is associated with pyrethroids, using AS-PCR techniques.

Insecticide susceptibility test results for deltamethrin (0.05%) confirmed a mortality rate of 96%, which, according to WHO criteria, indicates “possible resistance” (mortality range ≥90% but <98%) [21]. This intermediate resistance level suggests that resistance is developing in the *Aedes aegypti* population in Nagapattinam district, despite deltamethrin continued relative effectiveness. This study corroborates earlier studies that have reported similar outcomes. For instance, a study conducted in Nigeria, reported a 95.09% mortality rate against deltamethrin, suggesting the development of resistance [22]. However, other studies have reported confirmed resistance to deltamethrin in *Aedes aegypti*, with mortality rate less than 90%. A study conducted by [23] in Nouakchott, Mauritania, reported a significantly lower mortality rate of 36% when *Aedes aegypti* were exposed to a lower concentration of deltamethrin (0.03%). Similarly, another study conducted in Delhi, India, reported a high resistance status in *Aedes aegypti*, with mortality rates ranging from 64.4% to 74.3% against deltamethrin [24]. In contrast, research led by [25] in Ethiopia reported a 100% mortality rate against deltamethrin in *Aedes aegypti*, indicating full susceptibility to the insecticide. Consistent with this, [26] documented the high mortality rates in *Aedes aegypti* from colombia, further supporting the susceptibility to deltamethrin.

In addition, compared to deltamethrin, *Aedes aegypti* from Nagapattinam exhibited a stronger resistance to permethrin (0.75%), with a mortality rate of 84%, indicating confirmed resistance according to WHO criteria (WHO, 2022) (confirmed resistance <90%). A similar trend was observed in St. Andrew Jamaica, where *Aedes aegypti* population showed high resistance, with mortality rate ranging from 0% - 8%, interestingly, despite permethrin being in use for only the past two years in that region, resistance has already developed, suggesting a rapid selection pressure [27]. A parallel outcome was reported in Indonesia, exhibited less than >90% mortality rate in *Aedes aegypti* upon exposure to 0.75% permethrin [28]. Another investigation from the northern districts of West Bengal, India, reported a mortality range 60% to 83.3% in *Aedes aegypti* exposed to permethrin, further confirming the resistant status in various regions [29]. The observed variation in insecticide resistance status in the present study ranging from possible to confirm resistance against deltamethrin and permethrin differs from other findings across various regions that have been reported in *Aedes aegypti* populations. These discrepancies may be due to the geographic locations or variation in insecticide usage pattern. Considering the reduced susceptibility to pyrethroids observed in bioassays, molecular level investigation was carried out to detect the presence of kdr mutation, in particular V1016G mutation in domain II in the *vgsc* gene in *Aedes aegypti* population from the Nagapattinam district. The results from our study demonstrated the presence of the V1016G mutation in heterozygous mutation (V/G), among several mosquito samples that survived insecticide exposure. The detection of both 60bp (Valine allele) and 80bp (Glycine allele) products confirmed the heterozygous status of the mutation. V1016G mutation is a well-studied single amino acid substitution that confers resistance, particularly to pyrethroids [30]. However, this particular mutation has not been extensively studied in Tamil Nadu, therefore, there are few previous studies that support the present findings. Also studies have reported the presence of homozygous mutation among the *Aedes* populations. For instance, a study by [31] indicated a higher prevalence of homozygous individual compared to heterozygous in *Aedes aegypti* population from Thailand. Interestingly, findings from Thailand have documented the presence of multiple *kdr* mutation-including S989P, V1016G, and F1534C, occurring as triple heterozygotes (S/P989+V/G1016+F/C1534), which were found at high frequencies that are highly linked to pyrethroid such as both deltamethrin and permethrin [32]. Similarly, another research conducted in Srilanka, first reported the presence of V1016G mutation along with F1534C, and S989P in *Aedes aegypti* populations [33].

A study by [34] reported the presence of the V1016G mutation in *Aedes aegypti* from Makassar, Indonesia, attributing its emergence to prolonged use of the insecticides, and geographic isolation of islands like Sulawesi have led to local mutation events. In contrast, a study on Indian *Aedes aegypti* populations demonstrated the presence of the V1016G mutation, which was linked with the S989 but not with other major *kdr* mutation [35]. The current study provides the insecticide resistance status of *Aedes aegypti* population from Nagapattinam district, Tamil Nadu. The resistance status was further confirmed with molecular basis, aligning with similar findings reported worldwide. Further studies involving sequencing of the *vgsc* gene are warranted to better characterize the mutation pattern and their association with insecticide resistance in these population. To date, there is no available data on the presence of the V1016G *kdr* mutation in *Aedes aegypti* population from Nagapattinam district. Moreover, these findings highlight the need for continuous resistance monitoring and the implementation of integrated vector management measurements to overcome the risk of widespread resistance among the mosquito population.

## Conclusion

The present study has undertaken to assess the insecticide susceptibility status of *Aedes aegypti* mosquitoes collected from the various locations of Nagapattinam District and tested against widely accepted insecticides in the market worldwide: Deltamethrin and permethrin, both belongs to the class of Pyrethroids. The results from the WHO tube test revealed that the mosquito populations remained possible resistance to deltamethrin, whereas resistance to permethrin was evident from the sampling sites. This findings, indicates the evolving resistance patterns in the local populations and highlights the need for continuous monitoring to guide effective vector control strategies. To further investigate the molecular basis behind the observed resistance, resistant samples were subjected to allele-specific PCR targeting the V1016G knockdown resistance *(kdr)* mutation in the Voltage-gated sodium channels gene. Molecular analysis confirmed the presence of heterozygous form (V/G), located in the domain II of the *vgsc* gene .These findings emphasize the importance of integrating molecular diagnostics with routine insecticide resistance surveillance. Early identification of resistance markers like V1016G can inform timely vector control interventions and support the sustainable use of insecticides in disease-endemic regions like Nagapattinam district.

## Acknowledgements

The authors acknowledge the gratitude to the Indian Council of Medical Research (ICMR), Delhi, for research fellowships, and to the Department of Biotechnology at the Central University of Tamil Nadu for providing lab facilities and support. The authors thank Tata Institute for Genetics and Society (TIGS) for providing mosquito traps, consumables support.

---

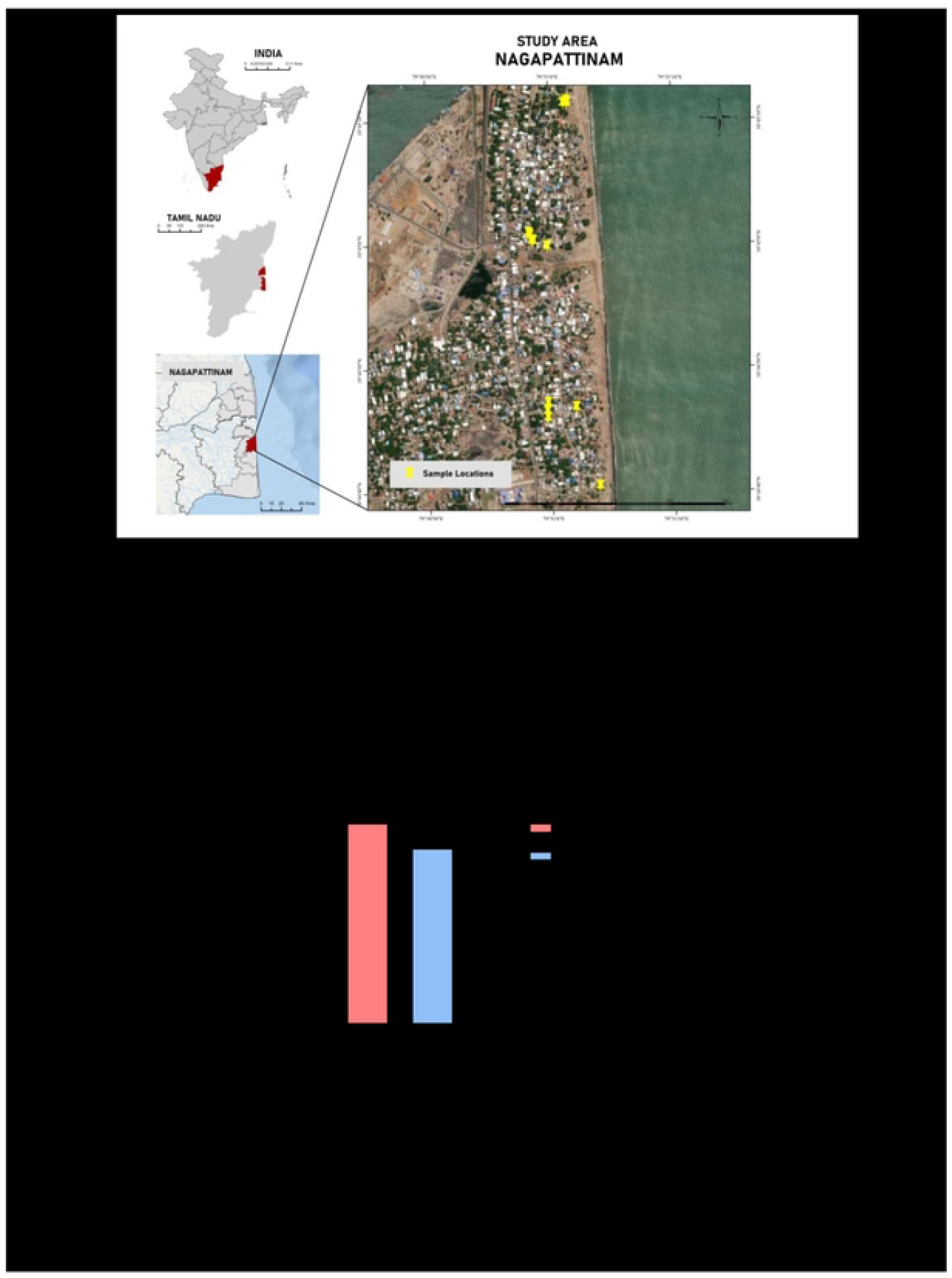

